# UPϕ phages, a new group of filamentous phages found in several members of *Enterobacteriales*

**DOI:** 10.1101/2019.12.27.889675

**Authors:** Jason W. Shapiro, Catherine Putonti

## Abstract

Filamentous phages establish chronic infections in their bacterial hosts, and new phages are secreted by infected bacteria for multiple generations, typically without causing host death. Often, these viruses integrate in their host’s genome by co-opting the host’s XerCD recombinase system. In several cases, these viruses also encode genes that increase bacterial virulence in plants and animals. Here, we describe a new filamentous phage, UPϕ901, which we originally found integrated in a clinical isolate of uropathogenic *Escherichia coli*. UPϕ901 and closely related phages can be found in published genomes of over 200 other bacteria, including strains of *Citrobacter koseri*, *Salmonella enterica*, *Yersinia enterocolitica*, and *Klebsiella pneumoniae*. Its closest relatives are consistently found in urine or in the blood and feces of patients with urinary tract infections. More distant relatives can be found in isolates from other environments, including sewage, water, soil, and contaminated food. Each of these phages, which we collectively call “UPϕ viruses,” also harbors two novel genes of unknown function.

## Introduction

Phages are often described by their potential to kill their hosts. Obligately lytic phages kill their hosts following infection, whereas temperate phages may lie dormant as prophages within lysogenized bacteria for several generations before entering a lytic cycle. While killing by phages has immediate consequences for bacterial ecology and has led to the revival of phage therapy for treating bacterial infections (Kortright et al 2019), many phages also carry genes that alter bacterial behavior (Warwick-Dugdale et al 2019; Mai-Prochnow et al 2015). The effects of these phage-encoded genes range from modifying photosynthesis in cyanobacteria (Sieradzki et al 2019) to producing toxins in potential pathogens (Waldor and Mekalanos 1996).

Filamentous phages in the family *Inoviridae* go beyond the standard dichotomy of viral lysis and lysogeny. Instead, the majority of characterized inoviruses maintain productive infections over multiple bacterial generations, without killing their hosts. In many cases, these phages integrate as tandem repeats into their hosts’ genomes at a locus called the *dif* site (Mai-Prochnow et al 2015). The *dif* site is a 28bp region at the bacterial terminus that includes two short palindromic regions recognized separately by the site-specific recombinase pair XerC and XerD. In bacteria with this system, XerCD is responsible for resolving chromosome dimers during DNA replication and cell division (Carnoy and Roten 2009). Filamentous phages co-opt this system by carrying their own copy of the *dif* site within their genomes, causing XerCD to confuse phage DNA for bacterial DNA. After integration, new phages are produced at relatively low rates and may be maintained indefinitely within the bacterial population (Val et al 2005).

These filamentous phages can have dramatic effects on their hosts’ phenotypes. In *Vibrio cholerae*, the phage CTXϕ encodes the toxin genes responsible for bacterial virulence in humans (Waldor and Mekalanos 1996). Similarly, filamentous phages have been associated with the virulence of other human pathogens (Gonzalez et al 2002) and agricultural pests (Yamada 2013; Kamiunten and Wakimoto 1982). In other cases, filamentous phages have been found to alter bacterial motility (Jian et al 2013) and biofilm formation (May et al 2011; Rice et al 2009). Despite their myriad effects on hosts, these phages remain understudied. As of this writing, there are only 45 recognized members of *Inoviridae* in RefSeq, and only 33 with established taxonomy by the International Committee on Taxonomy of Viruses (ICTV). Nonetheless, recent work has shown that thousands of potential inoviruses can be found as prophages in published bacterial genomes and metagenomes (Roux et al 2019).

Here, we describe a novel filamentous phage, UPϕ901, discovered as a prophage in a clinical *E. coli* isolate from a patient urine sample. UPϕ901 is most closely related to the non-integrating phages I2-2 and IKe in the *Lineavirus* genus of the *Inoviridae*. Like many filamentous phages, UPϕ901 integrates as a tandem repeat at the *dif* site in its hosts. Using the phage sequence from our isolate as a query, we searched for close relatives of UPϕ901 in published bacterial genomes and assemblies. We found closely related prophages in over two hundred strains of *E. coli*, *Citrobacter koseri*, *Klebsiella pneumoniae*, *Salmonella enterica*, and *Yersinia enterocolitica*. The most similar phages tended to come from patient urine samples or from the blood or feces of patients with urinary tract infections; more distant relatives can also be found in soil, water, animal feces, and contaminated food. We refer to the collection of related filamentous phages as “UPϕ viruses.” In several cases, identical UPϕ strains were found infecting multiple bacterial genera.

## Methods

### Bacterial strains in lab

Clinical isolates of *E. coli* and *C. koseri* (see Table 1) were provided by the Wolfe Lab at Loyola University Chicago and were originally isolated as part of separate studies on the urinary microbiota of women with and without symptoms of urinary tract infections or other urinary ailments (Garretto 2019, Price et al. 2016ab). These strains and their GenBank accessions are summarized in Table 1. *C. koseri* strains in this study have not been fully sequenced. *E. coli* JE-1 is an IncI plasmid-bearing strain typically used to propagate phage I2-2 and was obtained from the Felix d’Herelle Reference Center for Bacterial Viruses (Université Laval, QC, Canada).

**Table 1.** Bacterial strains in this study.

| Strain ID | Species | Has UPϕ? | WGS Accession |
| --- | --- | --- | --- |
| UMB0731 | *Escherichia coli* | No | NZ_RRWP000000000* |
| UMB0901 | *E. coli* | No | NZ_PKHH000000000 |
| UMB1220 | *E. coli* | Yes | NZ_RRVZ000000000 |
| UMB1526 | *E. coli* | Yes | NZ_RRVH000000000 |
| UMB5814 | *E. coli* | Yes | NZ_RRUX000000000 |
| UMB6655 | *E. coli* | No | NZ_RRUL000000000 |
| UMB6890 | *E. coli* | No | NZ_RRUI000000000 |
| UMB1389 | *Citrobacter koseri* | No | unsequenced** |
| UMB7451 | *C. koseri* | Yes | unsequenced |
| UMB8248 | *C. koseri* | No | unsequenced |
| JE-1 | *E. coli* | No | unavailable |

\* UMB0901 was sequenced as “E75” in its original BioProject (PRJNA316969)
\*\**Citrobacter* strains not sequenced at time of writing. Phage identified by PCR.

### Identifying phage in bacterial genomes

UPϕ901 was originally discovered as part of predicting phage sequences using PHASTER (Arndt et al 2016) in UMB0901 and other *E. coli* isolates in a previous study (Miller-Ensminger et al. 2018, Garretto 2019). In order to identify UPϕ901 in other published genomes, we took advantage of its integration at its hosts’ *dif* site as a tandem repeat. We defined a BLAST (Altschul et al 1990) query sequence for UPϕ901 as the region starting with the *dif* locus in UMB0901 and extending to the start of the first phage repeat. Throughout this paper, we will refer to the specific sequence (and homologs that are over 99% identical) as “UPϕ901.” More distantly related phages are collectively referred to as “UPϕ viruses.” The *dif* site itself is not repeated in full within the tandem duplications. We then used this single copy of the phage as a query for a blastn search using the NCBI web tool (https://blast.ncbi.nlm.nih.gov). This BLAST search identified hits to UPϕ901 in *C. koseri*, *E. coli*, *K. pneumoniae*, *S. enterica*, and *Y. enterocolitica*. Notably, neither PHASTER nor VirSorter (Roux et al 2015) consistently predicted UPϕ901 homologs in bacteria that had significant BLAST hits. This high false negative rate for these two tools is likely due to UPϕ901’s short genome and the reliance of these tools on information about known viruses. We have not tested the newer tool Inovirus_detector (Roux 2019) with genomes carrying UPϕ901 homologs.

We then downloaded all available and complete assemblies on NCBI for the five bacterial species listed above (as of August 2018). Each assembly was queried locally with tblastx to identify the specific start and stop positions of the putative phage regions. The start positions of each phage were confirmed by a separate BLAST search for the *dif* site specific to the bacterial host (Carnoy and Roten 2009), and stop positions were identified by confirming the first repeat in the BLAST results. Because not all assemblies correctly included phage repeats, some sequences did not include a repeat as an obvious stopping point. In these instances, we could only rely on the end position of the last significant hit in the query.

Last, we used searchsra (Levi et al 2018; www.searchsra.org) to check for UPϕ901 relatives in metagenomes. We then used the pileup.sh function from BBMap (Bushnell 2014) to check the completeness of UPϕ901 coverage for each putative searchsra hit.

### Phylogenetic analyses

We used anvi’o (Eren et al 2015) to facilitate gene clustering and annotation for the putative phage regions found in GenBank assemblies. Standard annotation tools fail to identify inovirus genes accurately, so we then used BLAST with individual phage IKe genes to confirm each gene’s correct annotation. Full-length UPϕ901 and most of the putative phage regions have every gene found in IKe (as in Figure 2). We then built a phylogeny for this subset of complete genomes with FastTree (Price et al 2010), using the concatenated alignment of each UPϕ901 gene amino acid sequence (genes *I*, *II, III, IV, V*, *VI*, *VIII*, *h1*, and *h2*). Individual gene alignments were found using MAFFT (Katoh and Standley 2013). Individual gene trees were also built using FastTree. Genes *VII* and *IX* were excluded, because they consist of only about 30 amino acids and were not consistently identified as genes. Gene *X* was also excluded as it is contained entirely within gene *II*.

Trees were visualized using iTOL (Letunic and Bork 2019) and the ape package (Paradis and Schliep 2018) in R (R Core Team). Additional metadata for tree visualization were obtained from NCBI, EBI, or from the literature. Key metadata included the type of material sampled for isolating bacteria (e.g. soil, blood, urine, feces) and the more general source of that material (e.g. environmental, animal, human, food). The full metadata associated with each sample is provided in the data repository (see Data Availability). Nine strains had no available metadata from a publication or database. Twenty-two other strains without available metadata were from the “100K Pathogen Genomes Project” (BioProject PRJNA186441; Weimer 2017) and are identified as “100K Project” in figures.

We also generated a bacterial phylogeny for each of the phage hosts, using tblastx to identify each of the 24 single-copy universal genes in each genome (Lang et al 2013), MAFFT for alignments, and FastTree for producing the final tree.

### Phage presence in culture medium

UPϕ901-infected strains were grown in lysogeny broth (LB) at 37°C with moderate shaking overnight. The following morning, 1 mL of each culture was removed, centrifuged at 16,000g for 1 minute, and the supernatants were filtered through 0.2 μm cellulose acetate syringe filters. 80 μl of each filtrate were then treated with OPTIZYME DNase I (Fisher BioReagents BP81071) for 30 minutes, followed by heat inactivation with EDTA at 65°C for 10 minutes. This DNase step was included to remove any bacterial genomic DNA from the supernatant, which could result from cell death due to lysis from other prophages or shearing forces. DNase-treated samples were incubated at 95°C for 10 minutes to degrade phage protein coats and expose the phage ssDNA. We then amplified phage genomic DNA using UPϕ901-specific primers, UPphi_shortFW (GGGTTTATCAGAGGGGTCAG) and UPphi_shortRV (AGGATGGCTCTAAGTCAACG). 16S PCR (63F/1387R primers) was used to confirm the absence of bacterial genomic DNA in the DNase-treated filtrates. The UPϕ901 and 16S PCRs were performed on 1 μl of unfiltered culture as positive controls.

### Phage infection assays

We tested the potential for UPϕ901 to infect new hosts using standard plaque assays by spotting 5 μl of filtrate from UMB0901 (as described above) on 0.7% agar LB overlays containing candidate bacterial hosts (UMB0731, UMB6655, UMB6890, JE-1).

We also tested for new infections using PCR. Colonies of prospective host strains were added to 5 mL of LB supplemented with 50 μl of filtered supernatant from UMB0901. Uninfected controls of each strain were also grown. These cultures were incubated for 18 hours. Following growth, 2 μl of each culture were used in a PCR designed to test for integration into the new host’s chromosome. The forward primer (IntCheck_FW: GTGTGTGGATGTGAATGGTG) in the assay was based on a conserved sequence in all tested hosts just upstream of the *dif* site, and the reverse primer (IntCheck_RV: CTGGCAGAACGAACGATTAC) recognized a region early in the phage genome. This PCR can only amplify DNA if UPϕ901 integrates into a new genome. An overnight culture of UMB0901 was used as a positive control for each reaction, and the uninfected cultures were used as negative controls.

### Data Availability

Data from this work is available at figshare (https://figshare.com/s/235048b35ae0617dac68).

## Results

### Initial identification

We originally identified UPϕ901 in a clinical isolate of *E. coli* (UMB0901) as part of separate work examining prophages present in the urinary microbiome (Garretto 2019). We then found nearly identical phages in three other patient isolates in our lab collection (UMB1220, UMB1526, and UMB5814). As with many other filamentous phages, UPϕ901 is integrated as a tandem repeat at the *dif* site in each of these genomes. In each case, we confirmed by PCR that the phage is shed from infected bacteria, as centrifuged culture supernatant treated with DNase is positive for UPϕ901 DNA (Figure 1a) but negative for bacterial genomic DNA (Figure 1b).

**Figure 1.**
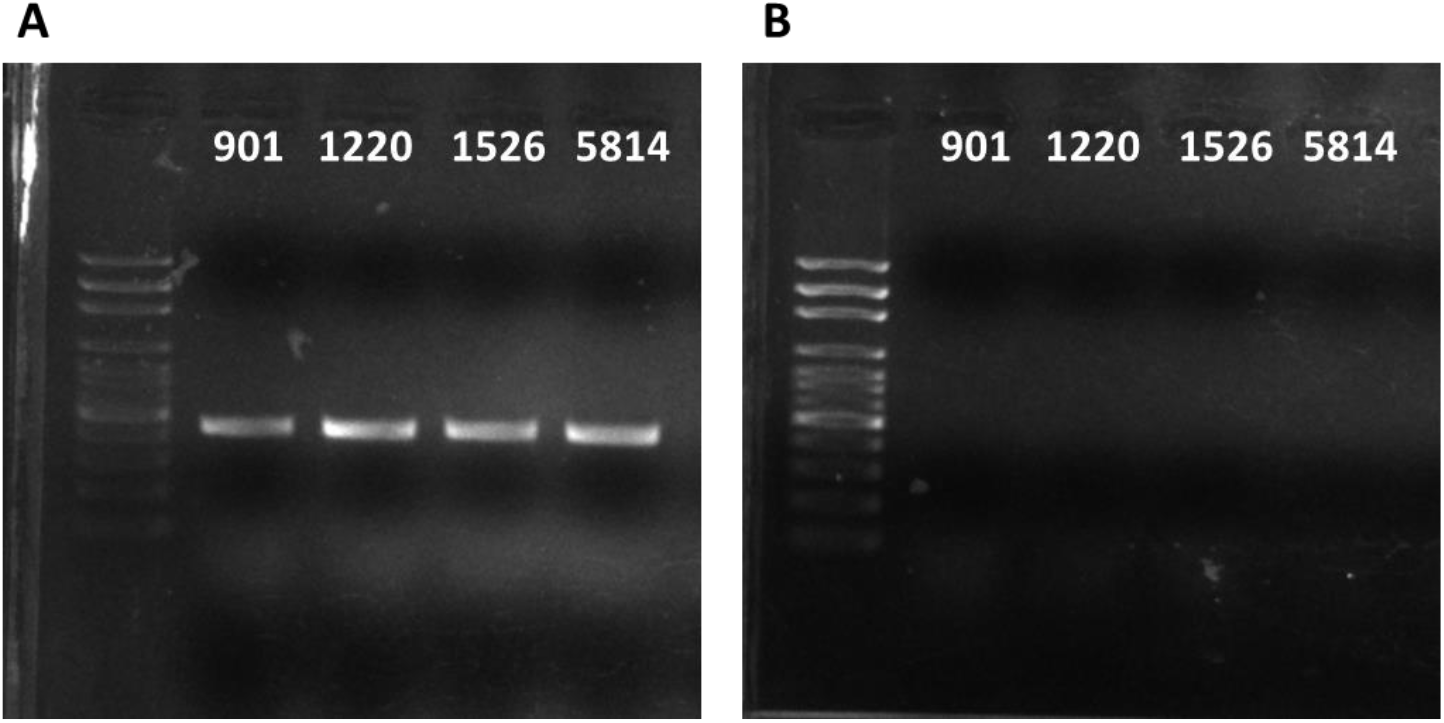
Phage DNA confirmation. PCR results using (A) UPϕ or (B) 16S primers after DNase treatment to remove bacterial genomic DNA from filtrates of UMB0901, UMB1220, UMB1526, and UMB5814 cultures.

### Comparative genomics

UPϕ901 is approximately 50% identical to phages IKe and I2-2 (70% coverage and ~70% amino acid identity for covered genes), each a non-integrating member of the *Lineavirus* genus of filamentous phages. These phages are also closely related to the F-specific filamentous phage, M13. Figure 2 shows how UPϕ901’s genome compares with these, with each gene shaded according to its amino acid similarity with the homolog in UPϕ901. UPϕ901 contains each of the “core” genes characteristic of filamentous coliphages with gene order preserved.

**Figure 2.**
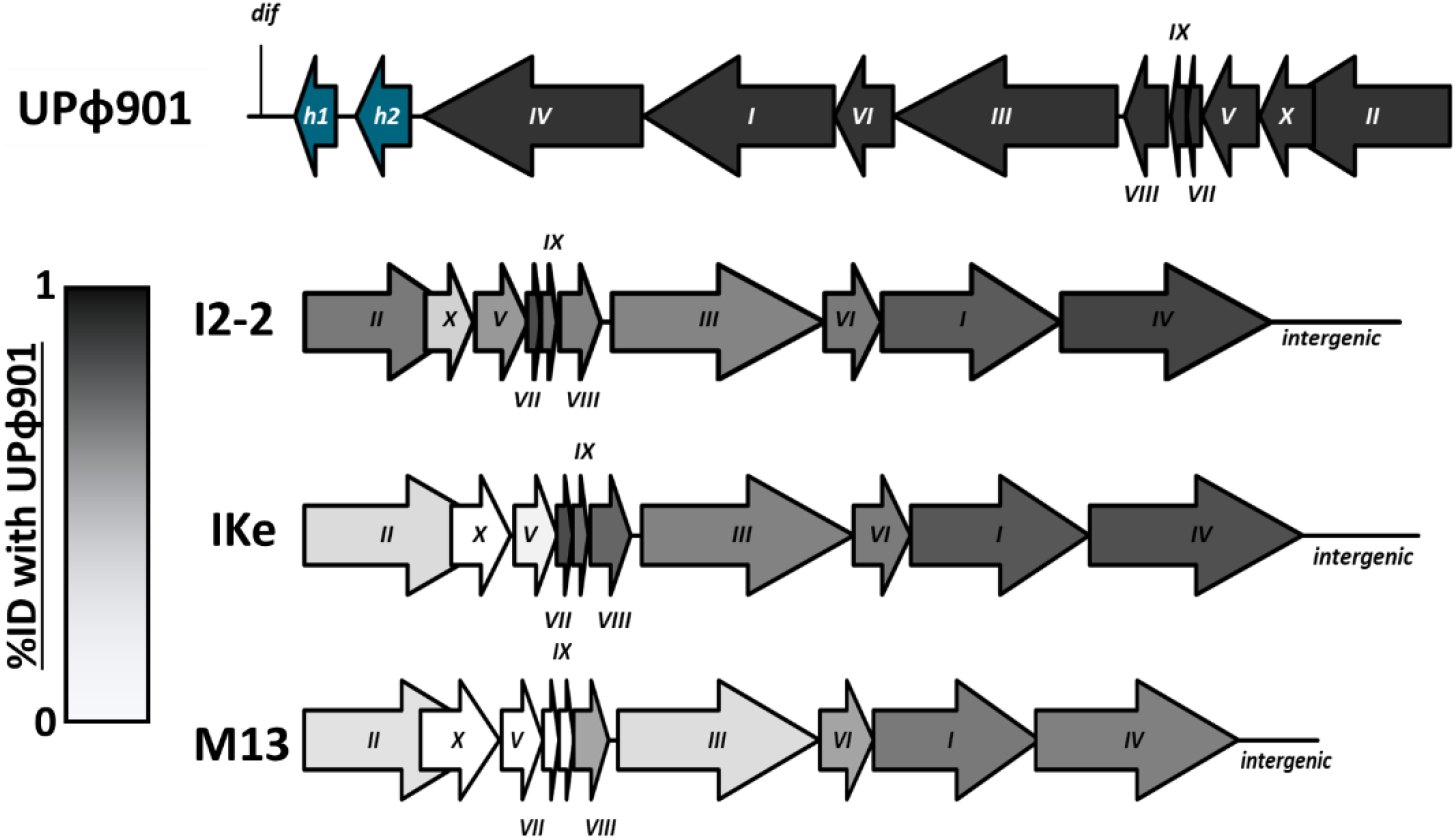
Comparison of UPϕ901 to related inoviruses. Roman numerals are the names for core inovirus genes common in M13 and the *Lineavirus* members. Genes shaded grey are darker if they are more similar to the UPϕ901 variant. ‘h1’ and ‘h2’ are two hypothetical proteins unique to UPϕ phages. The UPϕ901 genes are in reverse orientation due to integration within the host genome at the *dif* site.

Two notable differences set UPϕ901 apart from I2-2, IKe, and M13: 1) UPϕ901 includes a copy of the *E. coli dif* site at the start of its genome, enabling it to integrate via the XerCD recombinase system; 2) UPϕ901 encodes two genes of unknown function within a region that is intergenic in I2-2, IKe, and M13. These genes are not homologous to any known protein, and all attempts to find homologs of these genes in GenBank returned hits to prophages related to UPϕ901.

UPϕ901 strains are nearly 100% identical across infected clinical isolates of *E. coli* in our lab collection. The only distinction is that the attachment protein, g3p (encoded by gene *III*), in UPϕ901 has an additional repeat of a glycine-rich motif (“GGGES”) than the viruses in UMB1220, UMB1526, and UMB5814.

Using the UPϕ901 sequence as a query, we searched GenBank for homologous prophages in other bacteria. Closely related viruses could be found in published genomes of *E. coli*, *S. enterica*, *C. koseri*, *K. pneumoniae*, and *Y. enterocolitica*. We then downloaded all available assemblies from GenBank of these five species (26,397 in total) and used tblastx to identify related viruses integrated at *dif* sites. In all, 331 genomes harbored a similar prophage (see Table 2 for a summary by host). 229 of the 331 putative phages in GenBank assemblies contained each gene found in UPϕ901. For the remainder of this paper, we will refer to this collection of strains as “UPϕ viruses.” The 102 remaining genomes contained a significant hit to UPϕ901, but assembly or sequencing quality was inadequate to predict full phage genomes reliably. We then built a phylogeny for the 229 complete phages (Figure 3a). For each of the genomes in the phylogeny, we identified metadata (where available) for the sample source material (e.g. urine, feces, blood) and source environment (e.g. human, animal, environmental) and added these data to the tree visualization.

**Table 2.** Prevalence of UPϕ in GenBank Assemblies.

| Host Species | GenBank Assemblies | Assemblies with UPϕ |
| --- | --- | --- |
| *Escherichia coli* | 12,401 | 189 |
| *Salmonella enterica* | 9,144 | 95 |
| *Klebsiella pneumoniae* | 4,647 | 29 |
| *Yersinia enterocolitica* | 177 | 10 |
| *Citrobacter koseri* | 28 | 8 |

**Figure 3.**
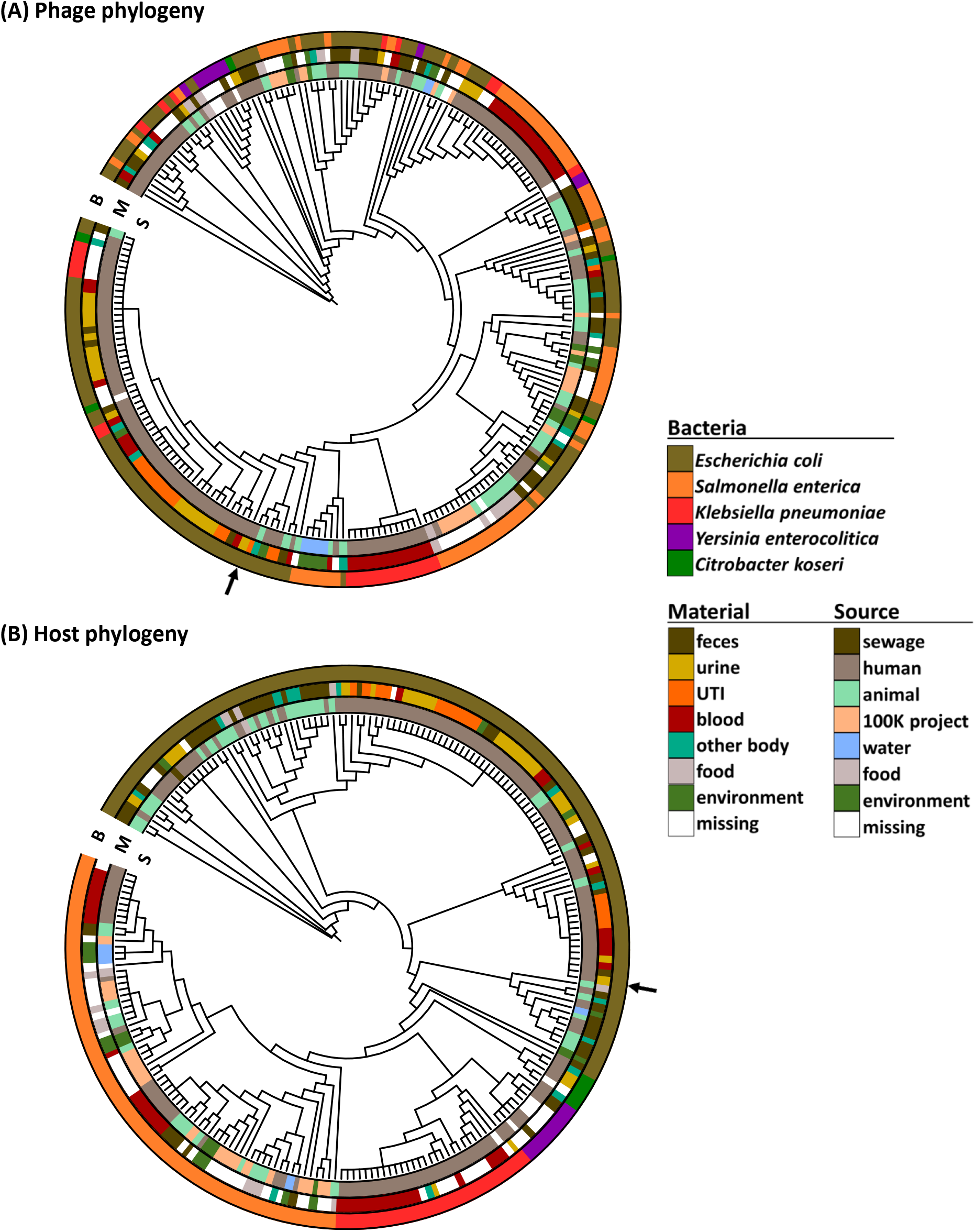
Phylogenies of UPϕ-related prophages in GenBank assemblies (A) and their hosts (B). The outer ring is the host bacterial species, the middle ring is the source material, and the innermost ring is the source environment. The black arrows indicate UPϕ901 (A) and UMB0901 (B).

UPϕ901 is indicated by an arrow in Figure 3a and is part of a clade of closely related phages found predominantly in *E. coli* from urine. Two large polytomies make up the majority of this group, with one consisting mostly of phages found in urine, and the other found in the blood or urine of patients with a urinary tract infection (UTI). Additionally, while most viruses in this urine-associated clade were found in *E. coli*, there are several cases where identical phages were also found within *C. koseri* and *K. pneumoniae*. This group also contains three samples from animals. The rest of the tree is dominated by phages found in *S. enterica* from both animals and humans, though phages infecting other bacteria are also present. Most of these remaining strains were found in feces or contaminated food. This portion of the tree also includes more distantly related phages found in *Y. enterocolitica*.

We also constructed a phylogeny for the host bacteria, using a common set of conserved single-copy genes (Lang et al 2013). The bacterial tree shows the expected breakdown by host species, with additional structure corresponding to sample material and whether the strain was from human or animal sources.

Individual phage gene trees (Supplemental Figure 1) show relatively low variation across the phages, with most genes composed of a small number of variants (Figure 4). Gene *III* stands out as having the most variation, with 95 distinct sequences among the 229 genomes. Examining the alignment for gene *III*, most of the variation is in the number of glycine-rich repeats in the region that also varied in our lab isolates.

**Figure 4.**
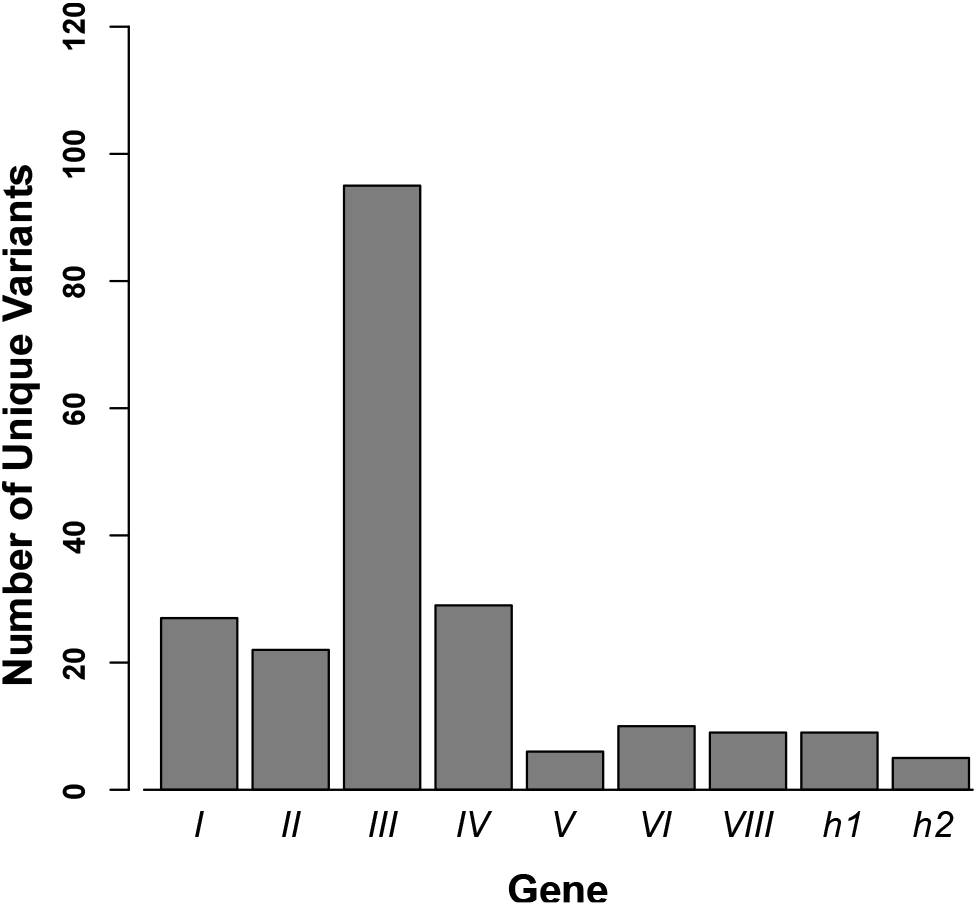
Number of unique sequence variants for each UPϕ gene used to build trees.

We also took advantage of the online tool searchsra (Levi et al 2018) to find relatives of UPϕ901 in metagenomes. Significant hits were returned for 4819 datasets, with samples from soil, gut, and aquatic environments. Of these, at least 1766 have over 95% coverage of the UPϕ901 genome.

### Identifying additional infections in the lab

Though only 28 *C. koseri* genomes were available, nearly one third carried a UPϕ virus. In two cases, these *C. koseri* phages were identical to prophage sequences found in urine *E. coli*s. Given this high frequency of infection and similarity to UPϕ901, we tested three unsequenced *C. koseri* isolates (UMB1389, UMB7451, UMB8248) using UPϕ901 PCR primers. The PCR results identified an integrated phage in UMB7451, but it was not actively produced, in contrast to the infected *E. coli* isolates (Supplemental Figure 2).

We also attempted to establish new infections using filtrate from UMB0901 cultures. Filamentous phages typically rely on conjugative pili as a primary receptor for infection (Mai-Prochnow et al 2015), and each *E. coli* carrying UPϕ901 in our lab collection also harbored an IncI conjugative plasmid. We identified three uninfected *E. coli* strains (UMB0731, UMB6655, and UMB6890) as candidates for testing UPϕ901’s ability to establish new infections. These strains all clustered together with UPϕ901-infected strains (UMB1220, UMB1526, UMB5814) in a gene presence-absence analysis of strains in our collection of urinary *E. coli* (Garretto 2019), contained an IncI plasmid, and were not already infected by UPϕ901.

Despite their potential as candidate hosts, UMB0901 filtrates that were confirmed to contain the phage did not produce plaques on any of these strains. We were also unable to observe plaques on *E. coli* JE-1, an IncI-bearing strain that is the standard host for UPϕ901’s closest known relative, I2-2 (Bradley et al 1983). Last, we used PCR to test for phage integration in these candidate hosts but were again unable to confirm new infections.

## Discussion

We have introduced a new filamentous phage, UPϕ901, found in multiple environments, with nearly identical strains found in clinical isolates of *E. coli* from urine. In addition to infecting *E. coli*, related viruses can be found in *S. enterica*, *K. pneumoniae*, *C. koseri*, and *Y. enterocolitica*. In several cases, identical phage genomes were found in multiple genera in similar environments. These phages are distinguished from characterized filamentous phages by the addition of two novel genes of unknown function. We have tentatively termed this collection of phages carrying these genes as “UPϕ viruses.”

These UPϕ viruses are most similar to the characterized phages IKe and I2-2 in the *Lineavirus* genus of *Inoviridae*, but IKe and I2-2 cannot integrate into the host genome and lack the accessory genes that appear to be unique to UPϕ phages. Given the prevalence of *dif* site integration among filamentous phages and their high sequence similarity to much of the UPϕ901 genome, it is likely that IKe and I2-2 evolved from an integrating ancestor.

Recent work has called into question the current taxonomy of filamentous phages and has suggested that the *Inoviridae* may require substantial revision into multiple new families of viruses (Roux et al. 2019). We are, therefore, hesitant to claim that UPϕ viruses deserve to be identified as a new phage genus. For the time being, we propose that the UPϕ viruses should be considered members of the existing *Lineavirus* genus, with the current members (IKe and I2-2) forming a subgenus of phages that have lost the *dif* site and nearby accessory genes. We acknowledge, though, that this taxonomy could change as the *Inoviridae* are revised.

A good deal of additional work remains to understand the prevalence and role of UPϕ viruses in different microbial communities. First, future research will need to characterize the two novel genes, *h1* and *h2*. These genes are found in a region of the genome that often includes genes that alter host behavior or virulence (Mai-Prochnow et al 2015). In the prototypical integrating inovirus, CTXϕ, this region encodes the cholera toxin genes and their regulators (Waldor and Mekalanos 1996). It appears likely that these two new genes interact with one another, but they share no homology to any known gene, and there is little to hint at their possible functions.

Future work that includes long-read resequencing of infected hosts and phages released into culture medium would also help to correct any errors in inferring the phage end positions. While we have identified 331 putative phages from genome assemblies, it was not always possible to predict phage end positions or the number of tandem repeats. These issues make it difficult to assess the pangenome of these phages, as some might carry additional accessory genes not found in UPϕ901.

We also attempted to determine the host requirements for establishing new infections of UPϕ901. We were able to confirm by PCR that new virions are released into the culture medium by infected bacteria, but we were unsuccessful in our attempts to infect new hosts. Our failure to observe new infections may be the result of lacking appropriate host strains for testing or due to low efficiency of establishing new infections. At the same time, UPϕ901’s closest relatives, I2-2 and IKe, each infect *E. coli* harboring IncI conjugative plasmids, and each *E. coli* strain infected by UPϕ901 in our collection also contained an IncI plasmid. It, therefore, appears likely that at least some of these phages rely on IncI-encoded pili, and future work with additional strains and different growth conditions will test this possibility.

It is also possible that different UPϕ viruses rely on different conjugative pili to initiate new infections. The phage attachment protein, g3p, is highly variable across strains, and each of these variants might correspond to different host specificity. Most of this variation was observed in a glycine-rich repeat region that links N- and C-terminal domains of g3p. Previous work showed that this region can diversify in phage IKe in a few generations in the lab, but no effects on phage fitness were observed (Bruno and Bradbury 1997). Those experiments, however, did not test for changes in host range.

Despite these technical challenges, UPϕ viruses represent an exciting new group of filamentous phages. Many inoviruses play important roles in modifying bacterial pathogens within eukaryotic hosts. It remains to be seen if UPϕ viruses, though frequently found in urine, affect the frequency of urinary tract infections or other aspects of urinary health. It is possible that UPϕ phages might affect bacterial ecology within the urinary environment without changing bacterial virulence. Similarly, it is unknown if these phages play any role in *S. enterica* food contamination. Perhaps more important will be understanding how frequently the phages shift hosts within different environments and whether these new infections alter the phenotypes of multiple genera within a community.

## Acknowledgments

We are grateful to the Wolfe Lab for generously providing strains of *E. coli* and *C. koseri* used in this work and to Putonti Lab members for helpful comments on the manuscript. This work is supported by NSF 1661357 to C. Putonti.

## Notes

https://figshare.com/s/235048b35ae0617dac68

