## Supplemental Figure for "UPϕ phages, a new group of filamentous phages found in several members of *Enterobacteriales*"

**Supplemental Figure 1. UPϕ Gene Trees (labeled by gene number)**

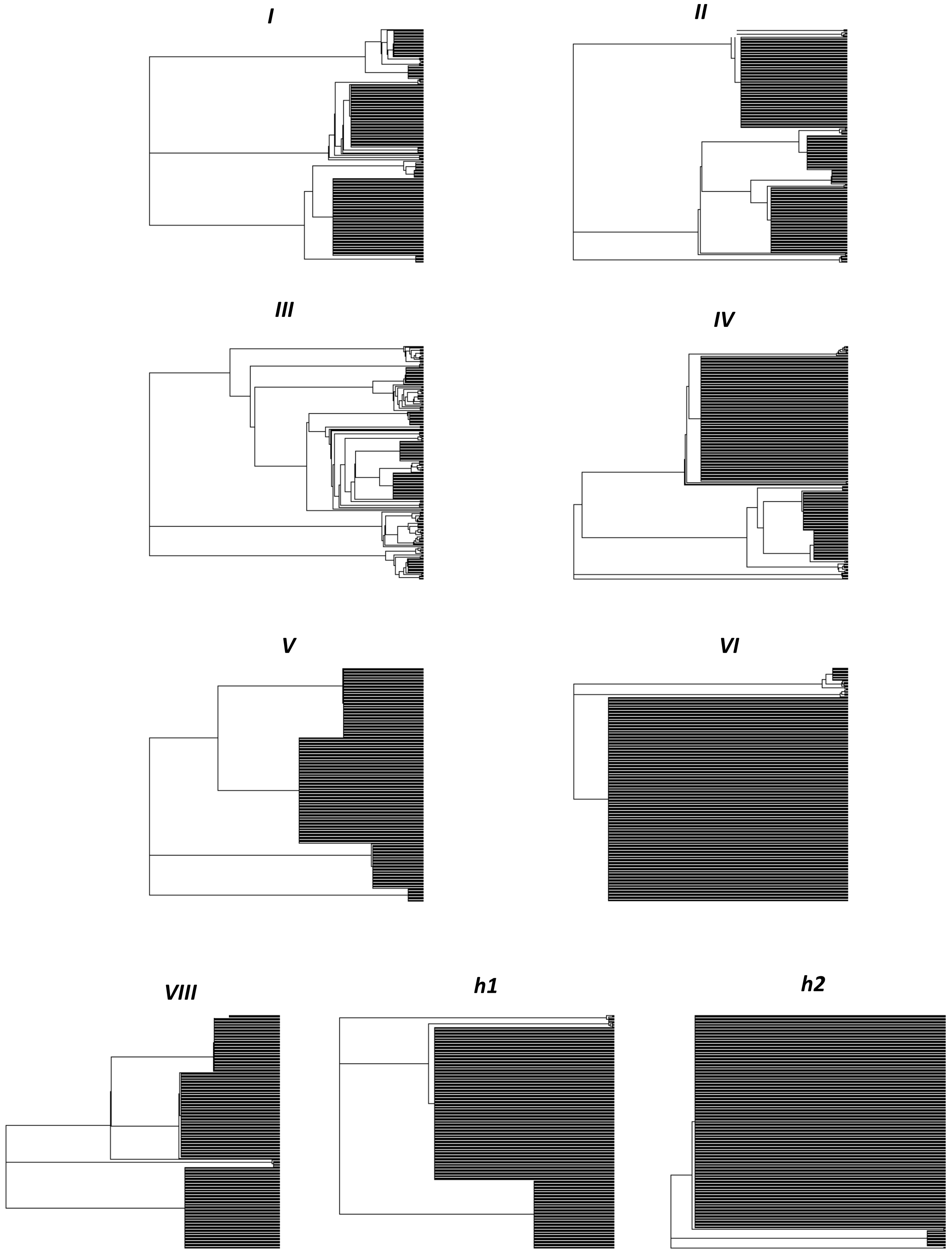

**Supplemental Figure 1. UPϕ gene trees**. Each tree made from MAFFT alignments and estimated using FastTree with visualizations using ape package in R. Branch lengths ignored to make the extent of polytomies more visible. Each tree labeled by the corresponding phage gene number.

**Supplemental Figure 2.** ***C. koseri*** **PCR Results**

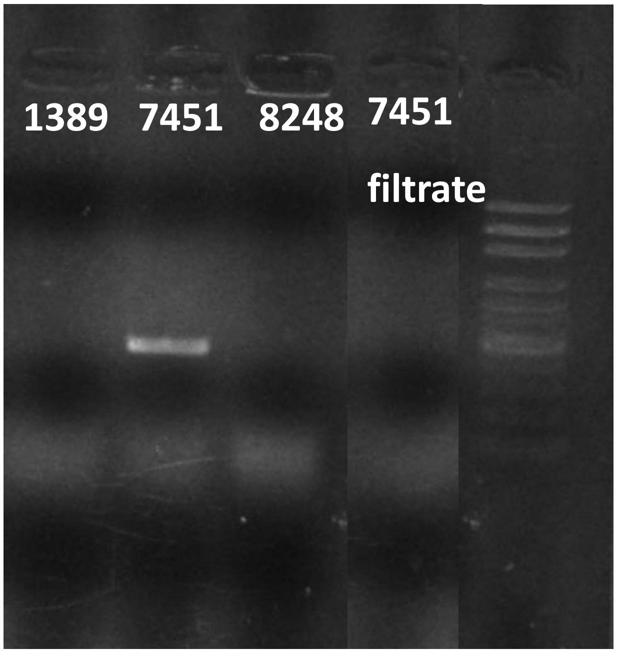


**Supplemental Figure 2.** ***C. koseri*** **PCRs.** A UPϕ virus was confirmed in UMB7451 but was not found in UMB1389 or UMB8248. It also did not appear to be secreted into the medium of a 7451 culture ("7451 filtrate"). Note that the column for "7451 filtrate" was from the same gel on the same day as the other results but was moved within the image (with no adjustments to scale or brightness) to improve the readability of results.
